# Adaptive divergence in shoot gravitropism creates hybrid sterility in an Australian wildflower

**DOI:** 10.1101/845354

**Authors:** Melanie J. Wilkinson, Federico Roda, Greg M. Walter, Maddie E. James, Rick Nipper, Jessica Walsh, Scott L. Allen, Henry L. North, Christine A. Beveridge, Daniel Ortiz-Barrientos

**Affiliations:** The University of Queensland, School of Biological Sciences, St Lucia QLD 4072, Australia; Universidad Nacional de Colombia, Departamento de Biología, Bogotá, Colombia; Monash University, School of Biological Sciences, Clayton Vic 3800; Floragenex, Inc., 4640 SW Macadam Avenue, Suite 200F, Portland, OR 97239, USA; University of Cambridge, Department of Zoology, Downing St., Cambridge CB2 3EJ, UK

**Keywords:** Local adaptation, intrinsic reproductive isolation, hybrid sterility, speciation, natural selection

## Abstract

Natural selection is a significant driver of speciation. Yet it remains largely unknown whether local adaptation can drive speciation through the evolution of hybrid sterility between populations. Here, we show that adaptive divergence in shoot gravitropism, the ability of a plant’s shoot to bend upwards in response to the downward pull of gravity, contributes to the evolution of hybrid sterility in an Australian wildflower, *Senecio lautus*. We find that shoot gravitropism has evolved multiple times in association with plant height between adjacent populations inhabiting contrasting environments, suggesting that these traits have evolved by natural selection. We directly tested this prediction using a hybrid population subjected to eight rounds of recombination and three rounds of selection in the field. It revealed that shoot gravitropism responds to natural selection in the expected direction of the locally adapted population. This provided an ideal platform to test whether genetic differences in gravitropism contribute to hybrid sterility in *S. lautus*. Using this advanced hybrid population, we discovered that crossing individuals with extreme differences in gravitropism reduce their ability to produce seed by 21%, providing strong evidence that this adaptive trait is genetically correlated with hybrid sterility. Our results suggest that natural selection can drive the evolution of locally adaptive traits that also create hybrid sterility, thus indicating an evolutionary connection between local adaptation and the origin of new species.

**Significance statement:** New species originate as populations become reproductively isolated from one another. Despite recent progress in uncovering the genetic basis of reproductive isolation, it remains unclear whether intrinsic reproductive barriers, such as hybrid sterility, evolve as a by-product of local adaptation to contrasting environments or evolve through non-ecological processes, such as meiotic drive. Here, we show that differences in a plant’s response to the pull of gravity have repeatedly evolved amongst coastal populations of an Australian wildflower, thus implicating a role of natural selection in their evolution. We found a strong genetic correlation between variation in this adaptive trait and hybrid sterility, suggesting that intrinsic reproductive barriers contribute to the origin of new species as populations adapt to heterogeneous environments.

## Introduction

Ever since Darwin’s work on the origin of species by natural selection (1), researchers have sought to understand how natural selection creates reproductive barriers between populations (2). On one hand, many studies have established that adaptation to contrasting environments often reduces migrant and hybrid fitness in the wild, a process commonly known as extrinsic reproductive isolation (3). These extrinsic barriers to reproduction can dramatically reduce gene flow between populations (4, 5); however, they only act in the local environment of populations and are therefore susceptible to changes in environmental conditions. Consequently, it is unclear whether the evolution of extrinsic reproductive isolation alone can complete speciation (4). On the other hand, intrinsic reproductive barriers such as hybrid sterility or inviability can accumulate regardless of environmental change (6) and therefore are expected to be more stable over time and contribute more reliably to the completion of speciation (2). One way in which natural selection can simultaneously create both extrinsic and intrinsic reproductive isolation is when the beneficial mutations that drive local adaptation in each population also fail to interact properly between populations through negative epistasis (7) (i.e., Dobzhansky-Muller genetic incompatibilities). However, given the paucity of examples that directly link loci that contribute to both local adaptation and intrinsic reproductive isolation, we remain ignorant as to whether natural selection drives speciation through the concomitant evolution of extrinsic and intrinsic barriers between populations (8).

Two notable examples genetically link local adaptation and intrinsic reproductive isolation, one via pleiotropy and the other via tight genetic linkage. Selection for pathogen resistance genes in *Arabidopsis thaliana* results in a pleiotropic effect of hybrid necrosis, which dramatically lowers reproductive success (9). In contrast, the tight genetic linkage between alleles selected for copper tolerance and alleles that cause hybrid mortality in *Mimulu*s *guttatus* led to divergence between populations growing next to copper mines and those occupying typical *Mimulus* habitats (10). Although plant-pathogen coevolution and tight linkage between genes performing various functions (e.g., stress tolerance and seed development) is a powerful example of a genetic mechanism that could often drive species divergence (11), we need more genetic and ecological studies to further understand when natural selection drives the correlated evolution of local adaptation and intrinsic reproductive isolation (12). So far, most results on the genetics of intrinsic reproductive isolation suggest that several mechanisms are driving the evolution of hybrid sterility across taxa (see Presgraves (13) for a review), such as genetic conflict (e.g., the evolution of distorter genes and their suppressors) (14), and parental conflict (e.g., endosperm failure in plants) (15). However, it remains unclear whether or not the local environment has a key role in creating intrinsic reproductive isolation (see Fishman and Sweigart (16) for a review of the mechanisms), thus limiting our understanding of the mechanisms creating genetic correlations between local adaptation and intrinsic reproductive isolation.

Here, we introduce a novel system of study where we take advantage of the repeated evolution of divergent growth habits to study the contribution of local adaptation to the evolution of intrinsic reproductive barriers in an Australian wildflower, *Senecio lautus* (G. Forst. ex Willd) (17, 18). Adjacent populations exhibit erect or prostrate growth habits dependent on whether they inhabit the sand dune or rocky headlands, respectively. Previous population genetic studies in *S. lautus* found that different sets of genes related by similar functions were repeatedly differentiated between populations with these two contrasting growth habits (19, 20). One of these sets contained genes belonging to the auxin pathway, where genes related to the transport and regulation of this key plant hormone had repeatedly diverged between erect and prostrate populations. In many plants, auxin genes are also involved in creating variation in height (21, 22), branching (23, 24) and reproduction (e.g. pollen tube growth (25)). Given the observed divergence in height and branching, with little gene flow between erect and prostrate populations of *S. lautus* (26, 27), the evolution of auxin-related genes may explain the presence of both adaptation and reproductive isolation in *S. lautus*. Studies across the plant kingdom have established that the auxin hormone has a conserved function in governing the way in which plants orient themselves to light and gravity cues (28). For example, mutant surveys in *Arabidopsis* revealed that many auxin-related genes (with functions influencing the biosynthesis, transport or signaling of the auxin hormone) were required for shoot gravitropism - directional growth response of the shoot against the pull of gravity (23, 28-30). With this in mind, we reasoned that if divergence in auxin-related genes contributed to the evolution of local adaptation, shoot gravitropism would be divergent between adjacent erect and prostrate populations and would differentially respond to natural selection within their local environments. Finally, if local adaptation were driving the evolution of intrinsic reproductive isolation in *S. lautus*, crosses between hybrid individuals with extreme differences in an adaptive trait would be genetically incompatible and not produce seed.

We tested these three predictions on seven coastal population pairs of *S. lautus* (Fig. 1a, Fig. S1 and Table S1), where there is strong evidence of local adaptation between adjacent Dune and Headland populations (31-34). Populations inhabiting sand dunes (Dune hereafter) are usually erect, while populations growing on adjacent rocky headlands (Headland hereafter) are usually prostrate (Fig. 1b). Erect and prostrate growth habits can also be found in related populations from the alpine regions of Australia, with a prostrate population inhabiting an exposed alpine meadow and an erect population inhabiting a sheltered alpine gully (Fig. 1c). Dune populations are continually exposed to high temperatures, high solar radiation, low salinity, and low nutrient sand substrate, whereas Headland populations are exposed to high salinity, high nutrients, and powerful winds (32). These Dune and Headland ecotypes are genetically grouped into two major monophyletic clades based on geography (eastern and south-eastern Australia), and adjacent Dune and Headland populations are often sister taxa, suggesting that they have evolved their contrasting growth habits independently multiple times (17, 27, 31, 35).

**Fig. 1.**
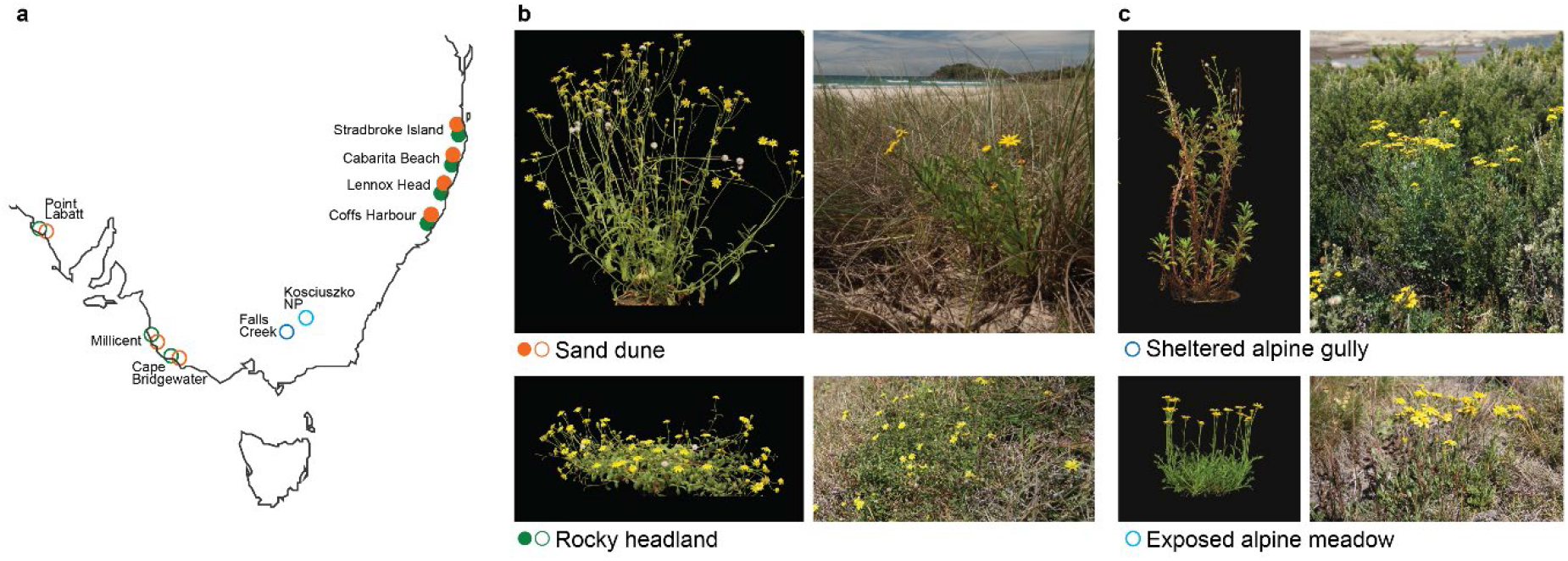
Sample locations and growth habit differences between adjacent *Senecio lautus* poopulations. **a**, Map of Australia showing locations of the 16 populations used in this study. The seven coastal localities have a Dune (orange) and Headland (green) population occurring adjacent to each other. The populations are split into two monophyletic clades (35), the eastern clade (closed cles) and the south-eastern clade (open circles). **b**, *Senecio lautus* native to the sand dunes have an erect growth habit and *S. lautus* native to the rocky headlands have a prostrate growth habit. **c**, Alpine populations of *S. lautus* include a sheltered alpine gully and an exposed alpine meadow, containing individuals with an erect and prostrate growth habit, respectively.

The two ecotypes also show variable levels of intrinsic reproductive isolation, where hybrid sterility in F1 Dune-Headland hybrids is generally weak. For example, there is <15% crossing failure at Lennox Head (31, 32) and <9% at Cabarita Beach (31). In contrast, hybrid sterility was found to be strong in F2 Dune-Headland hybrids from Lennox Head and an F2 generation created from four ecotypes (58% crossing failure (32, 36)). Genetic incompatibilities appear to largely be removed in the F2 generation as F3 hybrids are fertile (32, 36). Therefore, the inability to produce seed in F2’s is likely due to a small number of strong recessive negative epistatic interactions between these ecotypes (37, 38). Furthermore, demographic analyses recently revealed minimal gene flow levels between adjacent Dune and Headland populations (27), indicating that reproductive barriers (intrinsic or extrinsic) have prevented hybridization from occurring in the field. The presence of reproductive isolation between multiple locally adapted erect and prostrate populations of *S. lautus* provides an excellent opportunity to understand whether local adaptation driven by morphological and physiological traits can lead to the accumulation of intrinsic reproductive barriers.

## Results

### Divergence in gravitropic response is auxin dependent in *Senecio lautus*

To test the hypothesis that auxin-related genes drove the evolution of gravitropic differences between erect and prostrate *S. lautus* populations, we directly examined whether synthetic auxin and auxin transport inhibitors influence gravitropism differently between the ecotypes of a population pair. We grew Dune (n=90) and Headland (n=98) seeds collected from Lennox Head with synthetic auxin 2,4-D (2,4-Dichlorophenoxyacetic acid), and polar auxin transport inhibitor NPA (naphthylphthalamic acid) (39, 40). Because a gravitropic response requires an auxin concentration gradient, we reasoned that removing the gradient would reduce the gravitropic angle in Dune individuals (Fig. S2). As expected, the addition of synthetic auxin 2,4-D reduced the gravitropic angle more in Dune individuals than in Headland individuals (LR chi-square=18.49, P<0.0001). Similarly, the addition of auxin transport inhibitor NPA reduced the gravitropic angle in our experiments more in Dune individuals than in Headland individuals (LR chi-square=21.18, P<0.0001). This difference in hormone response between Dune and Headland individuals from Lennox Head, gives credence to our hypothesis that different gravitropic responses between erect and prostrate individuals reflect divergence in auxin-related genes.

### Repeated height and gravitropism divergence across *Senecio lautus* populations

To understand whether auxin-related genes might have repeatedly diverged with growth habit in *S. lautus*, we tested whether plant height (a simple growth habit trait) predicts shoot gravitropism in 16 *S. lautus* natural populations (Fig. 1a, Fig. S1 and Table S1). We measured the change in angle of a seedling’s shoot 24 hours after a 90° rotation (29, 41), where 0° is lack of a gravitropic response whereas 90° describes a complete re-orientation of the shoot and a large gravitropic response. We expected that short populations would have a smaller gravitropic angle than their adjacent tall population after rotation, if these traits were correlated. Across these populations, the average height of a population predicted its average gravitropic angle (Fig. 2a; F_1,13_=17.65, P=0.0010), with the average magnitude of gravitropism differing across the two monophyletic clades of this system (F_1,13_=32.58, P<0.0001). In addition, we found that plant height in natural environments (field) strongly correlates with estimates of plant height in common garden conditions (glasshouse) (F_1,9_=12.41, P=0.0065), indicating that differences in plant height are genetic and do not arise from plastic responses to the environment. Therefore, changes in gravitropism appear to be biologically correlated with divergent growth habit traits such as plant height, and this relationship has evolved independently in each clade.

**Fig. 2.**
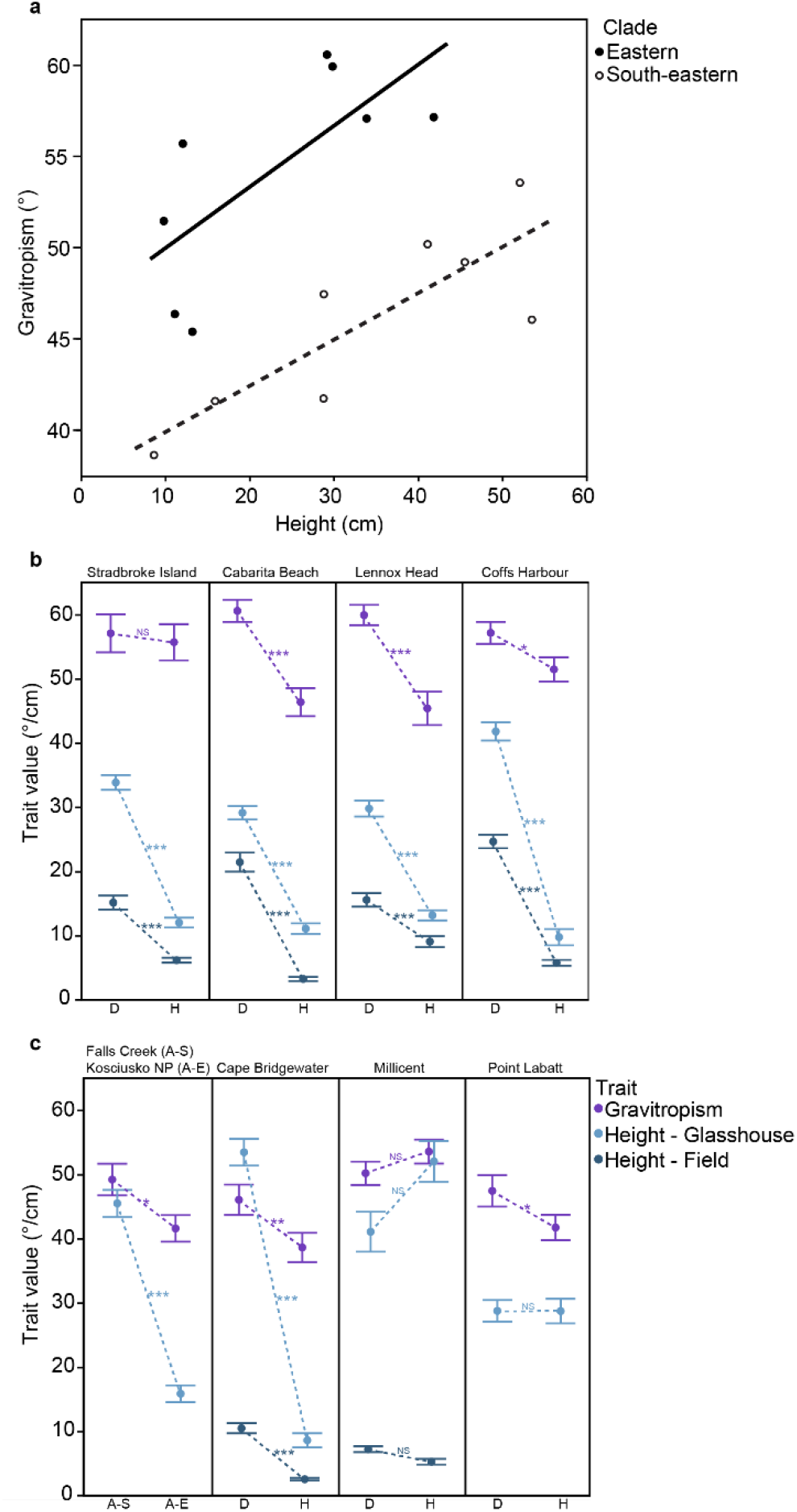
Gravitropism and height variation across 16 *Senecio lautus* populations. **a**, The correlation between gravitropism and height across *S. lautus* populations split into their monophyletic clades – see Fig. 1 for details. Each point in the graph represents a population mean where height was measured in the glasshouse and gravitropism was measured 24 hours after a 90° rotation. **b**,**c**, Divergence in gravitropism (°), height (cm) in the glasshouse and height in the field between adjacent *S. lautus* populations (D = Dune, H = Headland, A-S=Alpine Sheltered and A-E = Alpine Exposed). **b**, eastern clade and **c**, south-eastern clade. Height in the field for Falls Creek, Kosciusko NP and Point Labatt were not measured. Data are mean ± SE; one tailed Student’s t-test (Table S6), *P ≤ 0.05, **P ≤ 0.01, ***P ≤ 0.001, NS not significant.

We then investigated which of the seven adjacent Dune and Headland (and Alpine) population pairs drove the observed pattern. Four population pairs showed the expected correlation, where plants from the Headland population exhibited a smaller gravitropic angle and were shorter than plants from their adjacent Dune population (Fig. 2b, c). The expected pattern was also observed in divergent populations from the alpine region of Australia, where the exposed Alpine population in the meadow was shorter and had a smaller gravitropic angle than the population in the sheltered alpine gully (Fig. 2c). In the population pair at Millicent, plant height and gravitropism did not differ between Dune and Headland population pairs (Fig. 2c), possibly due to their similarity in environmental variables (35). At Stradbroke Island, we observed a difference in height in the expected direction but not gravitropism (Fig. 2b) and at Point Labatt, we observed a difference in gravitropism in the expected direction but not height (Fig. 2c), indicating that height and gravitropism are not always genetically correlated and alternate genes or pathways might be utilized (e.g., gibberellin controls dwarfism in some plants (42, 43)). Overall, these results are consistent with divergence in auxin-related genes evolving in parallel in coastal and alpine ecotypes in this system.

### Local adaptation drives the evolution of height and gravitropism

To directly assess the role of natural selection on the evolution of height and gravitropism, we conducted two independent sets of field adaptation experiments. In each field experiment, Dune and Headland parental and hybrid seeds were transplanted into replicated blocks at the sand dune and rocky headland at Lennox Head. In all field experiments, natural selection consistently favored the local Dune or Headland population over the foreign population (Fig. 3), indicating that our experiment was exposed to the positive selection that drove local adaptation in each environment.

**Fig. 3.**
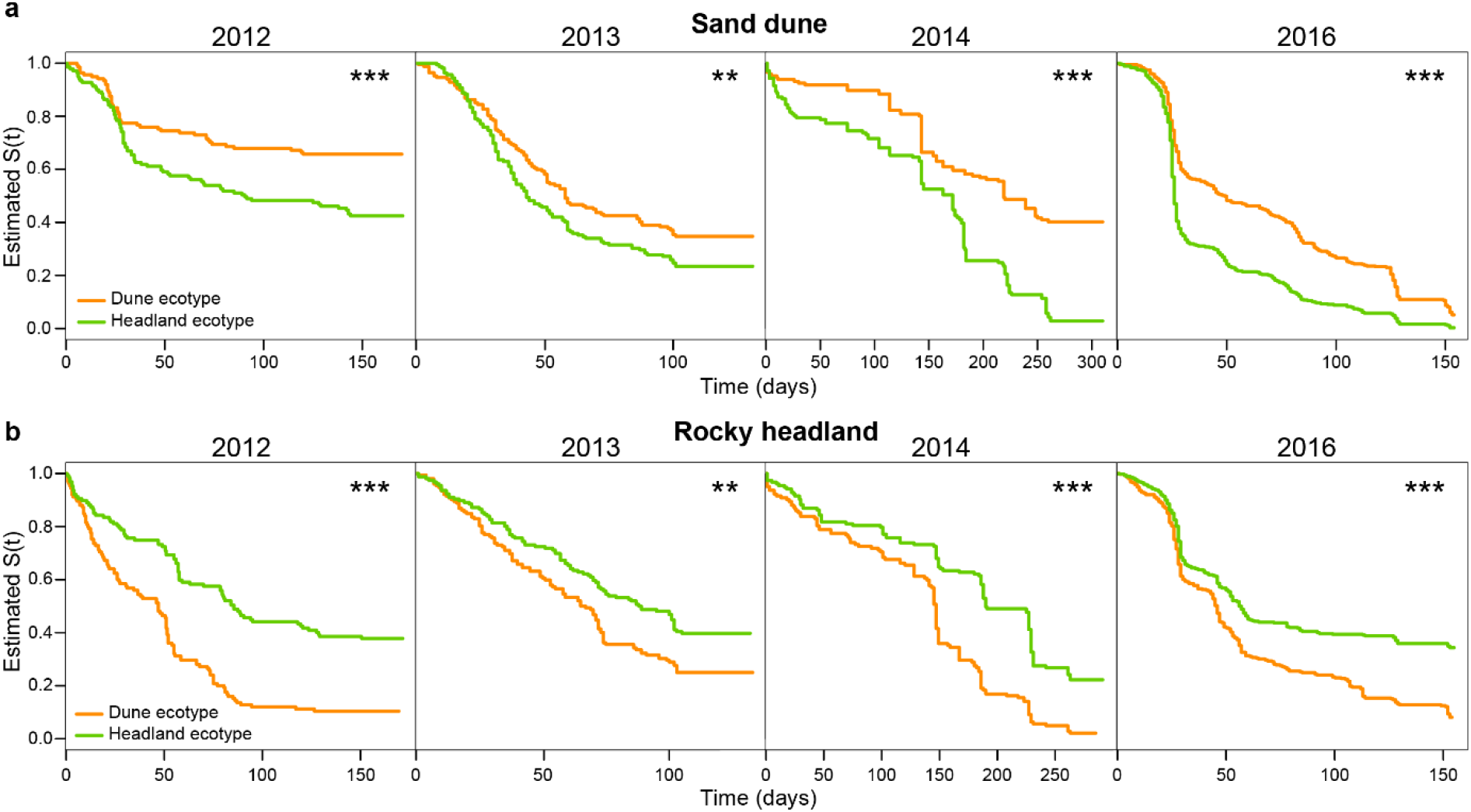
Parental survival curves in the height (2016) and gravitropism (2012-2014) adaptation experiments at the sand dune and rocky headland at Lennox Head. **a**,**b**, Survival (Estimated S(t)) over the length of the field experiments is shown for the Lennox Head Dune population (orange) and the Lennox Head Headland population (green) for four independent field selection experiments in the **a**, sand dune and **b**, rocky headland. Asterisks indicate a significant difference in mortality risk between the Dune and Headland ecotypes (**P ≤ 0.01, ***P ≤ 0.001).

First, we tested whether differences in height could drive differences in fitness in the rocky headlands. We hypothesized that if natural selection was driving the evolution of height, offspring produced by short hybrid parents would live longer than offspring produced by tall hybrid parents in the rocky headland. We focused on prostrate growth because it is likely the derived trait, given that the majority of *S. lautus* populations have an erect growth habit (19, 32). Our goal was to introgress Dune alleles associated with height onto a Headland genomic background to examine their effect on fitness in the headland environment. Briefly, we crossed Dune and Headland seeds from Lennox Head and then completed two rounds of backcrossing followed by one round of crossing between the tallest 10% and the shortest 10% (Table S2 and Fig. S3). We transplanted 558 of these seeds (from 28 families) into the rocky headland. As predicted, shorter hybrid parents produced offspring that lived longer in the rocky headland relative to offspring from taller hybrid parents (F_1,26.23_=4.87, P=0.0362). These results suggest that traits genetically correlated with plant height contributed to variation in early developmental fitness in the rocky headlands and contributed directly or indirectly to the evolution of divergent growth habit.

Next, we tested whether rapid adaptation to contrasting environments can lead to the evolution of gravitropic responses in the direction of the local population. We hypothesized that if natural selection was driving the evolution of shoot gravitropism, exposing an advanced recombinant population to multiple rounds of viability selection in the field would create genetic covariation between fitness and gravitropism. We crossed 23 Dune and 22 Headland individuals from Lennox Head to test this hypothesis and then maintained three independent genetic lines for eight generations (Table S3). This process disassembled allelic combinations and constructed an F8 recombinant hybrid population (26) (Fig. 4a). We planted 2,403 of these F8 seeds (from 89 families) into both the sand dune and rocky headland (Fig. 4b and Table S4) and conducted family-based truncation selection for three generations. The fittest families in each environment were selected based on the highest germination and survival (top 50%; see Methods for selection details). Siblings from these families were crossed (within a genetic line) to produce the next generation. In the F10 generation, we tested whether families with the largest number of survivors also produced offspring (F11) with the local gravitropic response under controlled conditions.

**Fig. 4.**
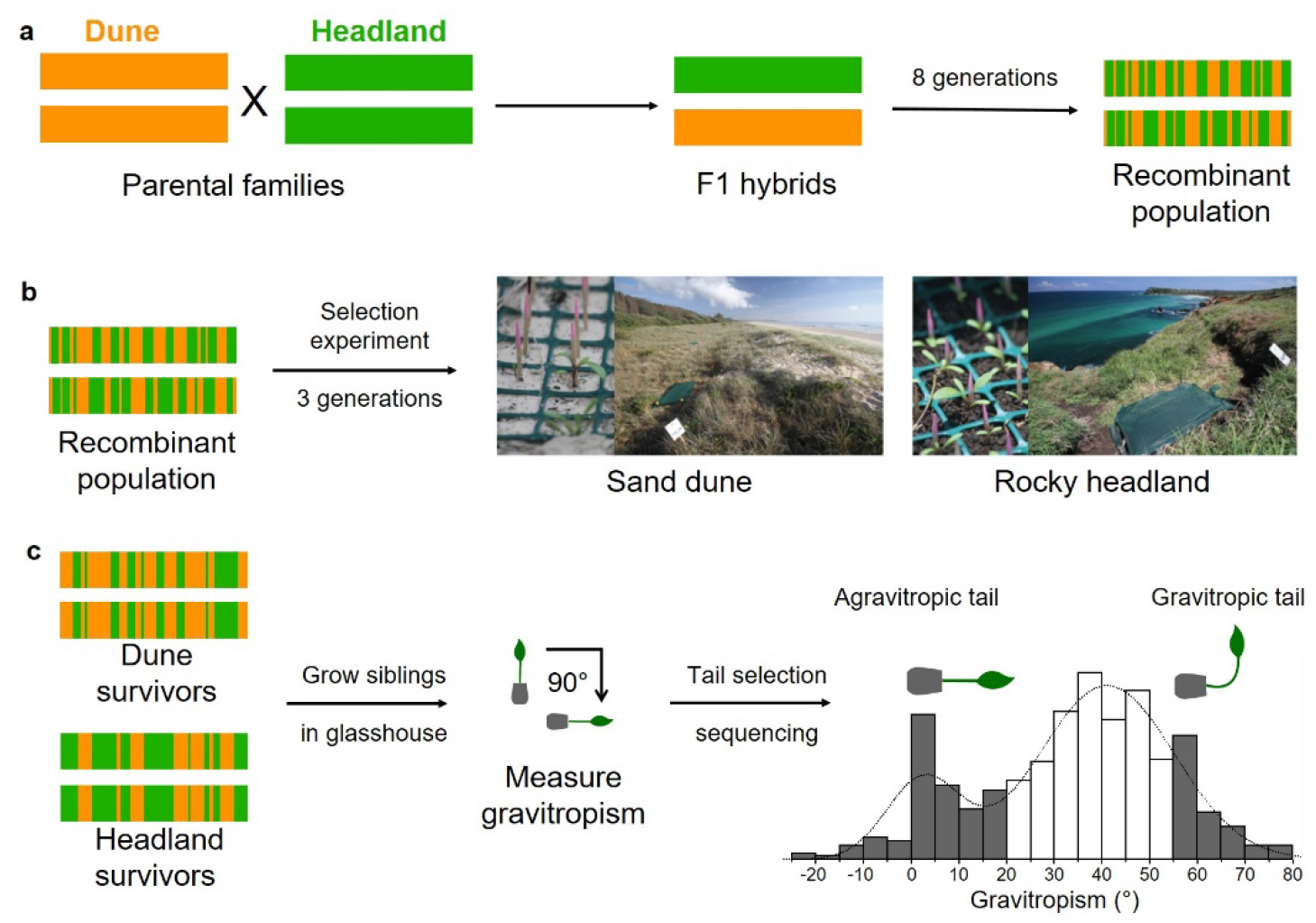
The creation of the recombinant hybrid generation, the design of the gravitropism adaptation experiments and sequencing of the tails of the gravitropic distribution. **a**, 23 parental Dune and 22 parental Headland individuals from Lennox Head were crossed randomly and with equal contribution for eight generations. **b**, Seeds from this F8 recombinant population were glued to toothpicks and transplanted into the sand dune and rocky headland at Lennox Head. Among-family based selection occurred for three generations (F8, F9 and F10), where full-siblings from the fittest families were grown in the glasshouse and crossed amongst their respective genetic lines (A, B and C) and their environment (Dune survivors or Headland survivors). An inbred control was kept in the glasshouse and underwent the same crossing scheme but free from viability selection. **c**, Gravitropism was measured in the F11 recombinant population by re-orientating the plant by 90°. Here, agravitropic plants are define as individuals with gravitropic angles <20°, while gravitropic plants have gravitropic angles >56° as they reorient their growth and subsequently grow upright. Individuals in the tails of the gravitropism distribution were sequenced on four lanes of the Illumina HiSeq 4000 platform.

In agreement with our prediction, F10 families with the largest number of survivors in the sand dune produced F11 offspring with a higher gravitropic angle (Table 1). We discovered that this relationship between fitness and gravitropism was driven by the fitness of the F10 dam family and not the F10 sire family (Table 1), suggesting maternal genetic effects might contribute to the evolution of gravitropism in the sand dunes. In contrast, we did not detect an association between the fitness of the F10 families and gravitropism of their offspring when we performed the experiment on the rocky headland. Instead, Headland survivors had a positive association between the number of days until death in a controlled environment (intrinsic viability) and gravitropism (Table 1), where individuals that died early in development had a smaller gravitropic angle (agravitropic). We are left with the conjecture that there could be intrinsic fitness costs for agravitropic alleles on a hybrid genomic background.

**Table 1.** General linear model for the effect of dam and sire on gravitropism (°) after a field selection experiment on a recombinant hybrid Dune and Headland population. Field selection experiments were performed on F8, F9 and F10 recombinant hybrid generations to achieve three rounds of selection in the sand dune and rocky headland at Lennox Head (see Fig. 4 for the experimental design). Dam and sire fitness are the F10 family fitness values for the individuals that were crossed to create the F11 offspring where gravitropism was measured. Intrinsic viability is the number of days until death of the F11 generation in the controlled temperature room. This experiment was conducted three times (temporal block) with three independent genetic lines.

| Source | Dune |  |  |  | Headland |  |  |  |
| --- | --- | --- | --- | --- | --- | --- | --- | --- |
|  | DF | SS | F-Ratio | P-value | DF | SS | F-Ratio | P-value |
| **Dam family fitness** | 6 | 8515.77 | 4.779 | **<0.001** | 6 | 1884.31 | 0.701 | 0.650 |
| **Sire family fitness** | 6 | 1806.62 | 1.014 | 0.424 | 5 | 2315.36 | 1.033 | 0.405 |
| **Intrinsic viability** | 1 | 260.85 | 0.878 | 0.352 | 1 | 5209.38 | 11.624 | **0.001** |
| **Genetic lines** | 3 | 1135.49 | 1.275 | 0.290 | 3 | 3357.19 | 2.497 | 0.067 |
| **Temporal block** | 2 | 193.14 | 0.325 | 0.724 | 2 | 2234.96 | 2.494 | 0.090 |

DF, degrees of freedom; SS, sum of squares.

We then tested whether adaptation to contrasting environments can recreate trait correlations observed in nature. As we have shown, there is a strong correlation between gravitropism and height across many (but not all) natural populations of *S. lautus*. We expect to lose this trait correlation in creating the hybrid population if different genes control the traits. If natural selection for these traits were strong, we could reconstruct the correlation after several rounds of selection in the coastal environments of the Dune and Headland populations. There was no genetic correlation between height and gravitropism in the control population grown only in the glasshouse (F_1,114.3_=0.08, P=0.7801, r^2^=0.04) indicating genes contributing to these traits are different and not pleiotropic. As predicted, the genetic correlation between height and gravitropism was strong after three rounds of selection in the rocky headland (F_1,169.5_=7.09, P=0.0085, r^2^=0.27) and weak after selection in the sand dune (F_1,151.3_=3.20, P=0.0756, r^2^=0.09). Together, these results indicate that natural selection can act on standing genetic variation and reconstitute genetic architectures favored in these coastal environments.

### The genetics underlying gravitropism in *Senecio lautus*

To better understand the genes underlying shoot gravitropism in *S. lautus*, we performed selective genotyping on the tails of the gravitropic distribution of the F11 hybrid populations (Fig. 4c; Dune survivors, Headland survivors and Control). We identified candidate gene regions containing the most extreme allelic differences between individuals with the smallest gravitropic angle (agravitropic tail, <20°; mean of tail = 6.46±1.10°, n=68) and the largest gravitropic angle (gravitropic tail, >56°; mean of tail = 62.03±0.45°, n=77). We found 55 sites (0.2% of all SNPs) across 49 genomic contigs (Data S1) with an allelic difference in the 99.8% quantile (0.15*<p*<0.22), indicating a polygenic basis for the phenotype. We discovered that these candidate gene regions disproportionally contained homologous genes with predicted gene ontology categories of transport and localization of molecules within and between cells (Table S5). This is consistent with expectations, as redistribution of auxin is required for a gravitropic response (28). Five of the 55 sites (11%) are located in gene homologs with functions related to the auxin pathway, including the second (*ENODL1*; early nodulin-like protein 1) (44, 45) and fourth (*ABA3*; molybdenum cofactor sulfurase) (46) most differentiated SNPs between the agravitropic and gravitropic F11 tails. *Arabidopsis wat1* mutants, an ortholog of *ENODL1*, are deficient in auxin production, display reduced auxin basipetal transport and have downregulated expression of many auxin-related genes, including those involved in response to auxin, auxin biosynthesis and transport (44, 45). *ABA3*, also named *LOS5* and *SIR3*, encodes a molybdenum cofactor sulfurase essential for the activity of several enzymes including for the plant hormone ABA and the auxinic compound sirtinol (47, 48). Genetic loss of function of *ABA3* causes impaired auxin signaling as well as reduced ABA levels (46).

Auxin-related genes, *ENODL1* and *ABA3*, have likely contributed to the adaptive and polygenic divergence of gravitropism in *S. lautus*. In both the *ENODL1* and *ABA3* genes, natural selection recreated the expected allelic frequency shift in the F11 tails towards the parent with the same trait (Data S1). For instance, the alleles favored in the agravitropic F11 tail were at high frequencies in the Headland natural population (*ENODL1* cytosine allele (C)=0.87 and *ABA3* guanine allele (G)=0.89), with the alternate alleles favored in the gravitropic F11 tail and the Dune natural population (*ENODL1* adenine allele (A)=0.69 and *ABA3* A=0.97). *ENODL1* and *ABA3* were in strong linkage disequilibrium in the survivors of the gravitropism adaptation experiment in the rocky headland (Fisher’s exact test, n=57, P=0.0008) and sand dune (Fisher’s exact test, n=48, P=0.0107), but not in the control population reared in the glasshouse (Fisher’s exact test, n=37, P=0.4093), suggesting natural selection has likely reconstituted this favorable allelic combination. Individuals with a *ENODL1* C/C and *ABA3* G/G genotype were associated with a reduction in gravitropism of 25° relative to all other genotype combinations (Fig. 5; t_34.30_=4.86, P<0.0001), indicating that not sensing the pull of gravity, or reacting to it, is a recessive trait. *ENODL1* and *ABA3* have gene homologs with functions in not only gravitropism and plant height but also in salt tolerance and pollen tube reception (44, 46, 49-53), suggesting that auxin-related genes could not only contribute to the adaptive evolution of gravitropism but also the evolution of reproductive trait differences between Dune and Headland populations that could affect seed production. Overall, our results suggest that divergence in auxin-regulated molecular processes contributed to the evolution of local adaptation to contrasting environments in coastal populations of *S. lautus*.

**Fig. 5.**
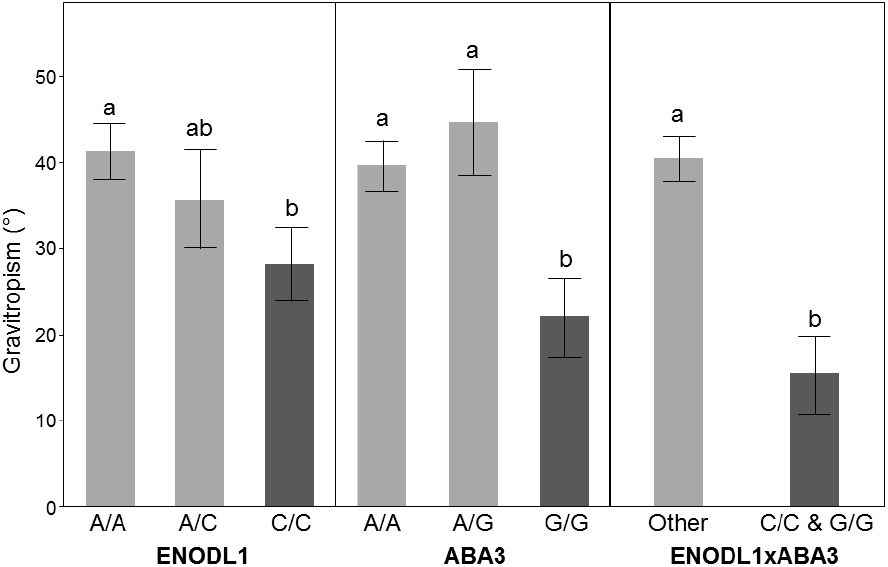
Association between *ENODL1* and *ABA3* alleles and gravitropism in *Senecio lautus*. The average gravitropism angle is shown for each allelic combination, independently and when the agravitropic alleles (dark grey) are combined. Different letters denote significant differences between genotypes at a significance level of α = 0.05.

### The consequences of gravitropism divergence for the evolution of hybrid sterility

To investigate whether divergence in auxin-related genes has consequences for reproductive compatibility between Dune and Headland populations, we measured crossing success between agravitropic and gravitropic F11 hybrids. Such crosses directly test whether a genetic correlation exists between gravitropism and hybrid sterility, and therefore evaluating the hypothesis that intrinsic reproductive isolation can evolve as a by-product of adaptation to local conditions. We therefore expected to observe increased hybrid sterility between F11 agravitropic and gravitropic individuals, and reproductive compatibility between individuals with similar gravitropism values. Hybrid sterility was defined as fewer than three seeds produced per flower head with at least three mating attempts for reciprocal crosses (fertile plants produce at least 30 seeds per cross per flower). We performed 28 crosses in families with an agravitropic response (within agravitropic tail, Fig. 4), 37 crosses in families with a gravitropic response (within gravitropic tail), and 67 crosses between these agravitropic and gravitropic families (between tails). Consistent with a genetic correlation between gravitropism and intrinsic reproductive isolation in *S. lautus*, we found that hybrid sterility was more common (Odd’s ratio=4.8x, P=0.0188) in crosses between F11 agravitropic and gravitropic plants (21%) than within each of these tails (5%; LR chi-square=6.86, P=0.0088). This pattern of crossing failures indicates that gravitropism alleles have linked or pleiotropic effects on hybrid sterility in *S. lautus*.

To assess what might be driving this association between gravitropism and hybrid sterility, we examined whether genetic line, F11 population, or specific individuals were correlated with hybrid sterility. Crosses within (n=77) and between (n=55) the three genetic lines, regardless of their gravitropic response, did not affect hybrid sterility (LR chi-square=1.10, P=0.2937). This result indicates that the hybrid sterility found in the F11’s is unlikely to be from genetic drift leading to incompatible differences between the genetic lines. Crosses within (n=80) and between (n=52) the three F11 populations (Dune survivors, Headland survivors and Control) did not affect hybrid sterility in agravitropic and gravitropic individuals (LR chi-square=0.15, P=0.6995), increasing the likelihood that gravitropism, and not another trait responding to selection in one of these environments, causes intrinsic reproductive isolation in these populations. Next, we determined whether specific F11 individuals (n=129) drove this association between gravitropism and hybrid sterility. Thirty-two individuals participated as one of the parents in a sterile cross; 26 of these individuals were crossed against a separate individual, and they successfully produced seeds. Thus, sterility is specific to each parental combination, consistent with the idea that hybrid sterility is polymorphic in the system, a result that echoes those found in other systems (54, 55). Finally, we found that crosses had symmetric effects on reducing hybrid fitness: only six crosses displayed differences in hybrid sterility in the reciprocal cross. Reciprocal effects on reproductive isolation are inconsistent with the contribution of maternal genotypes to survivorship in the field, suggesting that not only maternal genotypes contribute to sterility in these populations. Together, these results imply a genetic association between the adaptive divergence of gravitropism and hybrid sterility in *S. lautus*.

To investigate whether *ENODL1* and *ABA3* alleles were associated with hybrid sterility, we examined if the predicted allele counts of the F12 offspring from the F11 parents explained the proportion of failed crosses. We found that the interaction between *ENODL1* and *ABA3* did not have a significant effect on sterility (Table 2). The *ABA3* A allele, on the other hand, had positive effect on hybrid sterility, where individuals with the gravitropic favored A allele, which is dominant to the G allele, had a higher percentage of failed crosses (Table 2). This suggests that *ABA3* might genetically link adaptive evolution of shoot gravitropism with hybrid sterility in *S. lautus*. Overall, our results support the hypothesis that intrinsic reproductive isolation can evolve as a by-product of adaptation to local conditions.

**Table 2.** General linear model for the effect of *ENODL1* and *ABA3* alleles on hybrid sterility. The genotypes of the F12’s were predicted based on F11 parental genotypes with ambiguous genotypes removed. *ENODL1* is the allele counts for the *ENODL1* C allele in the F12’s, while *ABA3* is the allele counts for the *ABA3* G allele. *ENODL1* x *ABA3* is the effect of all observed allelic combinations between these two gravitropism candidate loci and the percentage of failed crosses.

| Source | DF | SS | F-Ratio | P-value |
| --- | --- | --- | --- | --- |
| *ENODL1* | 1 | 6.51 | 0.078 | 0.791 |
| *ABA3* | 1 | 925.52 | 11.076 | **0.021** |
| *ENODL1* x *ABA3* | 1 | 9.77 | 0.117 | 0.746 |

DF, degrees of freedom; SS, sum of squares.

## Discussion

Whether local adaptation commonly drives the formation of hybrid sterility and inviability is still a topic of debate (5, 16). These intrinsic reproductive barriers are believed to evolve late in the speciation process, often after extrinsic and prezygotic barriers have formed (e.g., immigrant inviability, or assortative mating) (4), and genetic drift or natural selection has led to the accumulation of genetic incompatibilities (2). There are few studies (9, 10) on the genetics of local adaptation and intrinsic reproductive isolation within a species, with most studies having focused on the effect of non-ecological evolution e.g., genetic conflict on the evolution of Dobzhansky-Muller incompatibilities (13). Our study is a step-forward in understanding the role of polygenic adaptation in creating hybrid sterility during the early stages of speciation. Here, we provide empirical evidence to suggest that natural selection is a major driving force behind the hybrid sterility found between recently derived erect and prostrate populations of *S. lautus*. Further, we identified variation in a trait in natural populations that could only result from divergence in specific hormonal pathways, thus suggesting the joint function of many genes could underlie the evolution of divergent traits correlated with reproductive barriers. The results from our set of experiments suggest a novel and broadly applicable explanation to the elusive link between the genetics of local adaptation and the evolution of intrinsic reproductive isolation.

Our results suggest that natural selection contributed to the colonization of extreme environments in *S. lautus*, like those found on rocky headlands along the coast. Previous population genetic results in this system (19, 26, 35) and those presented here suggest that the auxin pathway could facilitate plant adaptation to new habitats through concomitant changes in multiple developmental and architectural traits. Transitions from erect to prostrate growth and the associated traits of short stature and many branches, are common in plants that colonize coastal headlands (56-59), indicating that there are strong selective agents common to headland environments. For example, powerful winds or salty substrates could impose early selective pressures on traits controlled by auxins such as responses to mechanical cues (60) and halotropism (61-63). As the structure of hormonal pathways is generally conserved across plant species, the evolution of similar architectures in similar environments might prove to be a general mechanism to link adaptation with the incidental evolution of traits affecting species interactions such as ecological competition and reproductive success. For instance, in *Mimulus*, gibberellin (another essential plant hormone) has been recently implicated in local adaptation and associated with reproductive isolation between coastal and inland populations along the coast of California (42, 64). Like *S. lautus* Headland populations, coastal *Mimulus* populations are dwarf, salt-tolerant, and produce small amounts of gibberellin compared to inland populations (42, 65).

In *S. lautus*, previous results support a general link between local adaptation and intrinsic reproductive isolation. Walter, *et al*. (36) found that additive variance responsible for local adaptation in four ecotypes was lost in the creation of an F2 hybrid generation that displayed elevated levels of hybrid sterility. A similar occurrence of hybrid sterility occurred in the F2’s that created the F11 generation used here (32), suggesting that genetic incompatibilities are segregating in multiple populations and ecotypes of *S. lautus*. However, the other generations of the hybrid population (F3-F11) are highly fertile, implying that a large proportion of the incompatible alleles between Dune and Headland populations were removed in the F2 generation. Remarkably, we can recreate intrinsic reproductive isolation in the F11 hybrid population when we sort F11 families according to the strength of their gravitropic response; crosses between individuals with divergent gravitropism values have a reduced ability to produce seed. A likely explanation is that incompatibility alleles segregate at low frequencies in the population after the F2 generation, but we artificially increased the frequency of the incompatible alleles by selecting individuals with extreme gravitropism. Altogether these results provide strong evidence that adaptive traits, such as shoot gravitropism, are genetically correlated with hybrid sterility in *S. lautus*.

The mechanisms by which gravitropism and hybrid sterility are genetically correlated could be pleiotropy (e.g., Bomblies and Weigel (9)) or genetic linkage (e.g., Wright, Lloyd, Lowry, Macnair and Willis (10)), or a combination of both for polygenic traits. We speculate that pleiotropic effects of *ENODL1* and *ABA3* genes contributed to the correlated evolution of adaptive gravitropism and hybrid sterility. For example, *ABA3*, a strong candidate to generate variation in gravitropism, has been associated with pollen sterility in *Arabidopsis* (53), and it is known to contribute to salt tolerance in other organisms (51). It is not difficult to speculate evolutionary scenarios where hybrid sterility and gravitropism might be correlated responses to selection on saline environments: as the population adapts, compensatory mutations for reproduction can occur which are incompatible with genetic backgrounds from other populations. For *ENODL1*, we could not obtain many genotypes to properly test its role on hybrid sterility in *S. lautus*, but members of the *ENODL* family have a female-specific role in pollen tube reception (52). For example, in *Arabidopsis, enodl* mutants fail to arrest growth and rupture the entering pollen tube (52). The adaptive evolution of gravitropism through the maternal genotype in our adaptation experiments, also supports the notion that maternal *ENODLs* may be used to both communicate with the male pollen tube to enable reproduction and participate in adaptation to local conditions via its effects on gravitropism and correlated traits. The asymmetric reductions in seed set of *enodl* mutants contrasts with our symmetric reciprocal crosses. This discrepancy suggests that other factors other than maternal genetic effects contribute to reproductive isolation in *S. lautus*, possibly masking the effects of maternal genotypes in our experimental design. Future work will test the effect of *ENODL1* and *ABA3* on sterility using other genetic backgrounds and while considering novel genotypic combinations once we include other candidate genes in our genetic analyses.

Given that gravitropism has a polygenic basis and auxin related-genes have many pleiotropic effects on growth and development, and reproduction, our results suggest many genes likely drove local adaptation and only a subset of them contributes to hybrid sterility as they become part of a Dobzhansky-Muller incompatibility. This systems’ view of the evolutionary process, where pleiotropic effects are pervasive, could provide fertile ground for the origin of adaptations and new species. Future genetic studies focusing on isolating sets of loci causing hybrid sterility in *S. lautus* will reveal the molecular mechanisms by which intrinsic reproductive barriers evolve together with local adaptation and will allow testing of these novel predictions on the origin of new species. Overall, our study showcases a powerful strategy to explore the genetic basis of local adaptation and speciation in natural systems. We postulate that the evolution of hormonal pathways in plants provides a simple mechanism for the rapid modification of co-regulated traits that facilitate the colonization of new habitats and the correlated evolution of hybrid sterility.

## Materials and Methods

### Synthetic auxin and auxin transport inhibitor experiments

We tested if auxins govern gravitropism in *S. lautus* by evaluating gravitropic responses in seedlings treated with chemicals affecting auxin signaling. Specifically, we used 2,4-Dichlorophenoxyacetic acid (2,4-D), a carrier-dependent synthetic auxin (39, 66), and Naphthylphthalamic acid (NPA), an efflux inhibitor (40). Gravitropism was measured *in vitro* on agar plates on approximately 40 maternal families from Lennox Head sand dune and rocky headland. Seeds from an ecotype were combined in a falcon tube and were first sterilized with a quick rinse in 70% EtOH, followed by four 10-minute inversions in a sterilizing solution of 6% sodium hypochlorite and 1% Tween 20. Seeds were then rinsed three times with distilled water and vertically orientated on Murashiga and Skoog (MS) agar plates containing 0.15% MS, 0.05% 2-(*N*-morpholino)-ethanesulfonic acid (MES), 0.15% sucrose, 1% agar and 2,4-D and NPA at concentrations of either 0 µM, 0.5 µM, 5 µM, and 50µM. For the chemical concentrations, we used the following stock solutions: 2,4-D: 1mM in Ethanol and NPA: 10mM in dimethyl sulfoxide (DMSO). We created 1ml dilutions of stock solutions (in ethanol or DMSO), which were dissolved in 500 ml of media.

We placed eight seeds on each MS plate and incubated the plates at 21°C in a dark growth cabinet to avoid any light effects. After seven days, all plates were rotated clockwise by 90° and a photograph of each plate was taken 24 hours after rotation. All photographs were imported into ImageJ (67) to determine gravitropism by measuring the angle to which the stem reorientated to the horizontal (30, 68, 69). Seedlings were excluded from analyses if they were shorter than 5mm, contacted the plate’s edge, or germinated after rotation. We were left with a total of 188 seedlings to quantify the gravitropic response. The 50µM concentration of 2,4-D treatment was excluded, as less than six seeds germinated under this high concentration (Data S4). For each chemical, we used a mixed linear model using the lmer function of lme4 package in R v3.1.3 (70): y_ijkl_ = A_k_ + E_i_ + C_j_ + E_i_ x C_j_ + e_l(ijk)_, where agar plate (A_k_) is the MS plate that the seeds were grown on, ecotype (E_i_) is Dune or Headland and concentration (C_j_) are the 4 different concentrations of the chemical. Agar plate was fitted as a random effect, while ecotype, concentration and their interaction (E_i_ x C_j_) were fixed effects, and e_l(ijk)_ is the residual error. Gravitropism (y_ijkl_) was compared using a Type II Wald chi-square test.

### Phenotyping of natural populations

#### Height measurements

We measured plant height in all 16 populations in the glasshouse and 12 of the populations in their native field environment (Data S2 and Data S3). Height (vegetative) was measured as the vertical distance from the soil to the plant’s highest point that has vegetative leaves (flowers and stems are not included). In the field, we measured height in 32 individuals evenly across the range of each population. In the controlled conditions of the glasshouse, we sampled an average of 14 individuals per population after plants reached maturity.

For both the glasshouse and field measurements, we used a linear model to determine whether Dune populations were taller than their adjacent Headland pair: y_ijk_ = P_i_ + E_j(i)_ + e_k(ij)_, where pair (P_i_) is an adjacent Dune and Headland population at the same locality and ecotype (E_j(i)_) is Dune or Headland and is nested within pair. All factors are fixed effects and e_k(ij)_ is the residual error. Population height (y_ijk_) for each pair was compared using a one-tailed t-test (Table S6). The Alpine populations were also included with the prediction that the sheltered Alpine population (A03) would be taller than the exposed Alpine population (A07). All statistical results reported here were produced in JMP 13 (SAS 2015).

#### Gravitropism measurements

Gravitropism was measured *in vitro* on agar plates in a dark growth cabinet using seeds from all 16 natural populations. For each population, 2-4 seeds per family were grown for ∼40 maternal families (1,278 seeds in total). Plates were incubated, rotated, photographed, and gravitropism was measured in ImageJ (67), as outlined above. Overall, there was a 63.8% germination success, but seeds were excluded with the criteria above. This left a total of 736 seedlings across all 16 populations (57.6% of the total number of seeds planted; Data S2). To test the hypothesis that Dune populations would have a stronger gravitropic response in their stem than their adjacent Headland pair, we used a mixed linear model: y_ijkl_ = P_i_ + E_j(i)_ + A_k_ + e_l(ijk)_, where pair (P_i_) is an adjacent Dune and Headland population at the same locality, ecotype (E_j(i)_) is Dune or Headland and is nested within pair, and agar plate (A_k_) is the MS plate that the seeds were grown on. Agar plate was fitted as a random effect while the rest were fixed effects, and e_l(ijk)_ is the residual error. Gravitropism (y_ijkl_) measures were averaged for each population and compared between each population pair using a one-tailed t-test (Table S6). We then tested the correlation between height and gravitropism by performing a linear regression with mean height against mean gravitropism for all 16 populations, where populations were grouped into their respective clades (eastern and south-eastern). All statistical results reported here were produced in JMP 13 (SAS 2015).

### Field experiments

All field experiments were conducted at the sand dune and rocky headland at Lennox Head (NSW) in the exact location where native *S. lautus* grow. We tracked each individual by gluing each seed to a toothpick with non-drip superglue and placing them 1-2mm under the ground within a grid cell (Fig. 4) that was randomly assigned (for details see Walter, *et al*. (32)). 50% shade cloth was suspended 15cm above all plots to replicate the shade given by surrounding vegetation and was later replaced with very light bird netting. Seeds were watered twice a day to keep the soil moist and replicate ideal germination conditions to maximize the number of seeds in the experiment. Once germination plateaued for all genotypes watering was gradually ceased.

#### Height adaptation experiments

We created genetic lines that aimed to isolate height differences on a common genomic background (Fig. S3). Firstly, Lennox Head Dune and Headland seeds were grown and crossed to create an F1 generation. Secondly, we backcrossed to Headland parental plants for two generations to produce a BC2F1 generation. We then grew and crossed the tallest BC2F1 individuals among one another (n=16, tallest 10% of the population), and the shortest individuals among one another (n=18, shortest 10% of the population). See Table S2 for the number of individuals and families contributing to every generation to create these BC2F2 genetic lines. The BC2F2 seeds were planted into the rocky headland at Lennox Head in October 2016 (Australian spring). We planted five replicate plots, where each plot (1.08×0.33m) consisted of the same 12 families with four (occasionally three) individuals per family in each plot, totaling 1,116 BC2F2 seeds. Germination and mortality were recorded every day for 49 days, then every 3-4 days until day 79, and then weekly for the remainder of the experiment, until day 159 (Data S5).

We implemented a mixed linear model to test the hypothesis that individuals with short parents will have higher fitness in the headland environment: y_ijkl_ = H_i_ + F_j_ + B_k_ + e_l(ijk)_, where parental height (H_i_) was the average height of the parents measured in the glasshouse, family (F_j_) as individuals with the same parents, and block (B_k_) as the five replicate plots across the rocky headland. Parental height is a fixed effect, and family and block are random effects, and e_l(ijk)_ was the residual error. Offspring fitness (y_ijk_) was the total number of days alive in the rocky headland from planting. All statistical results reported here were produced in JMP 13 (SAS 2015).

#### Gravitropism adaptation experiments

We created an advanced recombinant population (F8) from 23 Dune and 22 Headland Lennox Head individuals using a North Carolina 2 breeding design (71, 72) as described in Roda, Walter, Nipper and Ortiz-Barrientos (26). Briefly, we replicated the construction of the F8 using three independent replicate crossing lines (A, B, and C), all derived from the same base population. See Table S3 for the number of families per replicate genetic line for every generation. F8 seeds were planted into the sand dune and rocky headlands in 2012 as described above. The fittest families (top 50%) within an environment (and genetic line) were selected using Aster modeling (73, 74) implemented with the ‘Aster’ package in R (70). The fitness components included germination and survival success, where germination was the total number of individuals that germinated in each family, and survival success was the total number of individuals per family that survived to day 85 in the F8 generation. One full sibling from the 101 selected families (50 Dune survivor and 51 Headland survivor families) and 44 families randomly selected from all F8 families for the control population were then grown in the glasshouse. Each individual was randomly assigned as a dam or sire and, after reaching maturity, crossed twice in a full-sibling, half-sibling crossing design. This approach maintained ∼100 full-sibling families for each population (Dune survivors, Headland survivors and Control). The same field experiment and selection procedure was conducted on the F9 (in 2013) and F10 generations (in 2014) for three rounds of selection, where survival success was measured at day 138 in the F9 generation, and at day 232 in the F10 generation. See Table S4 for the number of seeds and families planted per genetic line and environment for the three transplant experiments.

### Reference genome

Headland individuals from Lennox Head were used for the creation of an Illumina *S. lautus* reference genome. Firstly, we collected seeds from the Headland at Lennox Head and germinated seeds from two individuals by cutting 1mm off the micropyle side of the seed and placing in petri dishes with dampened filter paper. The seeds were placed in darkness for two days to allow for roots to grow and then transferred to a 12-hour light cycle in a constant temperature room at 25°C for seven days to allow for shoots to grow. Seedlings were then transferred into large individual pots with standard potting mix and grown in a glasshouse. These two individuals were crossed by rubbing flower heads together and collecting the seeds produced. Siblings from the seeds produced were grown and crossed by rubbing flower heads together to produce a family of individuals capable of self-fertilization (rare in *S. lautus*). One generation of selfing was completed to increase homozygosity. We extracted DNA from the leaf tissue of one individual using a modified CTAB protocol (75).

A draft genome of *S. lautus* was de novo assembled using second-generation short reads and AllPaths-LG 6.1.2 using default settings. We utilized a series of eight read libraries (Table S7). The reads were trimmed to remove residual adapter sequences and low-quality bases (minimum quality 15). The final assembly was ∼843 MB long and consisted of 96,372 scaffolds with an N50 of 21 KB. Although 843 MB is much shorter than the expected haploid size of 1.38 GB (76) of the whole genome, the BUSCO gene content completeness of 84% (5% fragmented and 11% missing) suggests that this assembly is primarily missing intergenic repetitive DNA sequences, which are notoriously difficult to assemble.

### F11 gravitropism measurements

In the F11 generation described above, we measured gravitropism as the angle of the stem after a 90° rotation of a seedling. This included 39 Dune survivor families, 37 Headland survivor families and 25 inbred control families with 12 individuals per family (1,212 seeds in total). These families were germinated in three separate batches ∼seven days apart. Briefly, we germinated the F11 seeds by cutting 1mm off the micropyle side of the seed and placing in petri dishes with dampened filter paper. The seeds were placed in darkness for two days to enable roots to grow and then transferred to light for 4 days for shoots to grow. Seedlings were then transferred into small square pots with standard potting mix in a constant temperature room at 25°C with a 12-hour light cycle. After one week of growing in the pot, the plants were rotated by 90° and a photograph of each individual was taken 12 hours after rotation. The photographs were imported into ImageJ (67) to measure gravitropism as described above. Data S6 contains gravitropism measures and dam and sire fitness values.

#### Gravitropism tests of selection

We implemented a linear model to test the hypothesis that high fitness Dune families will produce gravitropic offspring and high fitness Headland families produce agravitropic offspring. Independent models were used for the sand dune and rocky headland to test the effect of gravitropism on fitness in each environment: y_ijklmn_ = B_i_ + V_j_ + G_k(i)_ + D_l(ik)_ + S_m(ik)_ + e_n(ijklm)_, where temporal block (B_i_) is the three time points in which the F11 seeds were grown (∼seven days apart); intrinsic viability (V_j_) is the number of days until the death of F11 plants in controlled conditions; and genetic line, which consists of the three independent genetic lines (A, B and C), is nested within the temporal block (G_k(i)_). Dam fitness was nested in genetic line and temporal block (D_l(ik)_) and sire fitness was also nested in genetic line and temporal block (S_m(ik)_). Dam and sire fitness is the F10 family fitness values for the individuals that were crossed to create the F11 offspring where gravitropism was measured. All factors were included as fixed effects, and e_n(ijklm)_ was the residual error. Genetic line C was removed from analyses as it has little variation in fitness values, which means it did not converge. Shapiro-Wilk W test shows the residuals from the model are normally distributed for both the sand dune (W=0.98, P=0.3879) and rocky headland (W=0.98, P=0.2776). The linear model was performed in JMP 13 (SAS 2015).

#### Genetic association between height and gravitropism

We tested the genetic association between height and gravitropism after segregation in an advanced recombinant population. We implemented a mixed linear model for the three F11 populations (Dune survivors, Headland survivors, and a Control population) that accounts for family variation: y_ijk_ = H_i_ + F_j_ + e_k(ij)_, where gravitropism (y_ijk_) is the angle of the growth response 12 hours after a 90° rotation, height (H_i_) is the vertical distance from the soil to the top of the vegetative leaves, measured after maturity in the glasshouse and family (F_j_) is a random effect that consists of individuals that have the same dam and sire. The mixed linear model was performed in JMP 13 (SAS 2015).

#### Genotyping of F11 gravitropism tails

To isolate gravitropism candidate genes, we genotyped 77 gravitropic (>56°) and 68 agravitropic (<20°) F11 individuals (Data S6). We extracted DNA from leaf tissue using a modified CTAB protocol (75) and quantified the DNA using the PicoGreen reagent (Invitrogen, Carlsbad, CA). To determine which parent the alleles were derived from, we included 39 Dune parentals (D01) and 41 Headland parentals (H01). Leaves from the Lennox Head Dune and Headland natural populations were collected directly from the field and the same DNA extraction protocol was followed. Each F11 individual was duplicated in independent wells and libraries of Restriction-site Associated DNA (RAD) tags were created at Floragenex following Baird, *et al*. (77), but using the PstI restriction enzyme. We sequenced 380 samples on four lanes of an Illumina HiSeq 4000 with 91bp single-end reads at Floragenex. A total of 1.39 billion reads with a mean of 3.62 million reads per sample were produced. Reads were aligned to the Illumina reference genome using Bowtie 1.1.1 (78) and a FASTQ quality score of above 20. We then used SAMtools 0.1.16 (79) to create an mpileup file of all samples with a minimum Phred quality score of 10, minimum sequencing depth per sample of 6x and minimum per cent of population genotyped of 75%. This approach produced 26.8K variable positions (224K variants before filtering) with a minimum distance between variants of 50bp. The gravitropism candidate gene set consisted of SNPs in the 99.9% quantile of the distribution of differentiated SNPs between the gravitropism tails. The region of the scaffold containing the SNP was annotated using the BLASTx NCBI database (80).

#### Senecio lautus gravitropism candidate genes

We tested for overrepresentation of gene function and linkage disequilibrium between gravitropism loci in the gravitropism candidate gene set. A statistical overrepresentation test was performed in PANTHER (http://pantherdb.org/) using the TAIR identification for 32 unique gravitropism candidate genes matched to a reference list of 27,502 *Arabidopsis thaliana* genes. To calculate linkage disequilibrium between loci, a likelihood-ratio chi-square test was performed in JMP 13 (SAS 2015) with each F11 population independently (Dune survivors, Headland survivors and Control; Data S7).

### Intrinsic reproductive isolation

#### Gravitropism and hybrid sterility correlation

We tested whether hybrid sterility was associated with gravitropism by randomly crossing within and between the F11 gravitropic and agravitropic groups in a controlled temperature room and recording successful and failed crosses. To maximize sample size, all three replicate genetic lines (A, B, and C) were used across all three environments (Dune survivors, Headland survivors, and Control). A total of 138 crosses were completed (Data S8), 71 crosses between the tails and 67 within the tails (agravitropic tail = 29 and gravitropic tail = 38). Crosses were completed by rubbing multiple flower heads of two individuals and collecting the seeds produced from both plants. To remove genetic incompatibilities that might be caused by relatedness, crosses within the same family were not performed. Hybrid sterility (a failed cross) was considered when, in the presence of pollen, less than three seeds were produced per flower head from both plants (reciprocal crosses) with three mating attempts. Here, hybrid sterility could be caused by failure to produce fertilized ovules (prezygotic), or by fertilized ovules failing to develop into viable seeds (postzygotic). Six crosses displayed differences in sterility in the reciprocal cross (asymmetry), with one successful and one failed cross when using the same parents, and so were removed from the analysis (4 crosses between the tails and two within the tails).

If gravitropism contributed to divergence in *S. lautus*, we expected to observe an increase in hybrid sterility in crosses between gravitropic and agravitropic individuals. We performed a linear model in JMP 14 (SAS. 2018) to determine whether there was a significant association between cross-type (within vs between gravitropism tails) and hybrid sterility while accounting for genetic line and F11 population: y_ijkl_ = T_i_ + G_j_ + P_k_ + e_l(ijk)_, where tails (T_i_) is crosses within or between gravitropism tails, genetic line (G_j_) are the three independent genetic lines (A, B and C) and population (P_k_) are the three F11 populations (Dune survivors, Headland survivors, and Control). All factors were included as fixed effects, and e_l(ijk)_ was the residual error. Hybrid fertility (y_ijkl_) was compared using a likelihood-ratio chi-square test.

#### ENODL1 and ABA3 association with hybrid sterility

We implemented a linear model to test the hypothesis that *ENODL1* and *ABA3* alleles are correlated with hybrid sterility. From the F11 individuals that were crossed, we extracted their genotypes and predicted the genotypes of their offspring (F12). Genotypes that were ambiguous due to heterozygous parents were removed, which reduced the sample size to 61 crosses in which 11 were failed crosses (Data S9). We tested the effect of *ENODL1* and *ABA3* alleles and their interaction on the percentage of failed crosses: y_ijk_ = E_i_ + A_j_ + EA_ij_ + e_k(ij)_, where *ENODL1* (E_i_) is the allele counts for the *ENODL1* C allele in the F12’s; *ABA3* (A_j_) is the allele counts for the *ABA3* G allele in the F12’s; and *ENODL1* x *ABA3* (EA_ij_) is their interaction. All factors were included as fixed effects, and e_k(ij)_ was the residual error. Shapiro-Wilk W test shows the residuals from the model are normally distributed (W=0.94, P=0.6314). The linear model was performed in JMP 15 (SAS 2015).

## Acknowledgments

We are grateful to S. Smith, L.H. Rieseberg, M. Cooper, J. Engelstaedter, M.A.F. Noor and members of the Ortiz-Barrientos laboratory for insightful comments on previous versions of this manuscript. S. Karrenberg and S. Chenoweth provided instrumental feedback on M.J. Wilkinson Ph.D. dissertation. We thank P. Brewer for his help in the design and execution of gravitropism experiments. We would also like to thank everyone that helped with the field selection experiments: E. Johnston, E. Monley, G. Wilkinson, P. Wilkinson, A. Mather, S. Thang, K. Giarola, G. Lebbink, E. Firkins-Barriere, P. Arnold, J. Donohoe, B. Brittain, H. Liu, D. Bernal, M.C. Melo, T. Richards, J. Walter, L. Ambrose, B. Ayalon, S. Carrol, M. Gallo, A. Maynard, C. Palmer, and S. Edgley.

## Author Contributions

MJW, FR and DO conceived the projects and experiments. MJW conducted height adaptation experiments, reared and phenotyped various glasshouse populations and performed reproductive isolation experiment. FR and MJW performed physiological and molecular biology experiments. GMW reared and phenotyped the natural populations in the glasshouse with help from MJW and MEJ. GMW conducted the field experiment for the gravitropism adaptation experiments with assistance from MJW, MEJ, FR and DO. MJW, MEJ and HLN phenotyped natural populations. MEJ and MJW extracted the DNA from the F11’s and RN and JW called the genotypes. MEJ reared the plants and prepared the Illumina libraries for SA to assemble the *S. lautus* genome. CB guided the physiological experiments. MJW and DO wrote the paper with input from all authors. DO secured the funds and is mentor and supervisor for the research program.

## Notes

### Competing Interest Statement

The authors have declared no competing interest.

## References

1. C. Darwin, On the origins of species by means of natural selection (London: Murray, 1859), vol. 247, pp. 1859.

2. J. A. Coyne, H. A. Orr, Speciation (Sinauer Associates, Sunderland, MA, 2004).

3. H. D. Rundle, M. C. Whitlock, A genetic interpretation of ecologically dependent isolation. Evolution 55, 198–201 (2001).

4. O. Seehausen et al., Genomics and the origin of species. Nat. Rev. Genet. 15, 176–192 (2014).

5. E. Baack, M. C. Melo, L.H. Rieseberg, D. Ortiz-Barrientos, The origins of reproductive isolation in plants. New Phytol. 207, 968–984 (2015).

6. J. M. Coughlan, D. R. Matute, The importance of intrinsic postzygotic barriers throughout the speciation process. Philosophical Transactions of the Royal Society B-Biological Sciences 375 (2020).

7. D. Schluter, G. L. Conte, Genetics and ecological speciation. Proc. Natl. Acad. Sci. USA 106, 9955 (2009).

8. D. B. Lowry, R. C. Rockwood, J. H. Willis, Ecological reproductive isolation of coast and inland races of Mimulus guttatus. Evolution 62, 2196–2214 (2008).

9. K. Bomblies, D. Weigel, Hybrid necrosis: autoimmunity as a potential gene-flow barrier in plant species. Nat. Rev. Genet. 8, 382 (2007).

10. K. M. Wright, D. Lloyd, D. B. Lowry, M. R. Macnair, J. H. Willis, Indirect evolution of hybrid lethality due to linkage with selected locus in Mimulus guttatus. PLoS Biol. 11, e1001497 (2013).

11. M. Kirkpatrick, N. Barton, Chromosome inversions, local adaptation and speciation. Genetics 173, 419 (2006).

12. A. F. Agrawal, J. L. Feder, P. Nosil, Ecological Divergence and the Origins of Intrinsic Postmating Isolation with Gene Flow. International Journal of Ecology 2011, 435357 (2011).

13. D. C. Presgraves, The molecular evolutionary basis of species formation. Nat. Rev. Genet. 11, 175–180 (2010).

14. A. Burt, R. Trivers, Genes in Conflict: The Biology of Selfish Genetic Elements (Harvard University Press, 2006).

15. S. A. Johnston, T. P. M. den Nijs, S. J. Peloquin, R. E. Hanneman, The significance of genic balance to endosperm development in interspecific crosses. Theor. Appl. Genet. 57, 5–9 (1980).

16. L. Fishman, A. L. Sweigart, When two rights make a wrong: The evolutionary genetics of plant hybrid incompatibilities. Annu. Rev. Plant Biol. 10.1146/annurev-arplant-042817-040113 (2018).

17. I. J. Radford, R. D. Cousens, P. W. Michael, Morphological and genetic variation in the Senecio pinnatifolius complex: are variants worthy of taxonomic recognition? Aust. Syst. Bot. 17, 29–48 (2004).

18. I. Thompson, Taxonomic studies of Australian Senecio (Asteraceae): 5. The S. lautus/S. lautus complex., Muelleria (Royal Botanic Gardens, Melbourne, 2005).

19. F. Roda et al., Convergence and divergence during the adaptation to similar environments by an Australian groundsel. Evolution 67, 2515–2529 (2013).

20. M. E. James, M. J. Wilkinson, H. L. North, J. Engelstädter, D. Ortiz-Barrientos, A framework to quantify phenotypic and genotypic parallel evolution. bioRxiv 10.1101/2020.02.05.936450 (2020).

21. M. Xu, L. Zhu, H. Shou, P. Wu, A PIN1 family gene, OsPIN1, involved in auxin-dependent adventitious root emergence and tillering in rice. Plant Cell Physiol. 46, 1674–1681 (2005).

22. J. G. Wallace et al., Genome-wide association for plant height and flowering time across 15 tropical maize populations under managed drought stress and well-watered conditions in Sub-Saharan Africa. Crop Sci. 56, 2365–2378 (2016).

23. A. Gallavotti, The role of auxin in shaping shoot architecture. J. Exp. Bot. 64, 2593–2608 (2013).

24. M. a. Domagalska, O. Leyser, Signal integration in the control of shoot branching. Nat. Rev. Mol. Cell Biol. 12, 211–221 (2011).

25. J.-Z. Wu, Y. Lin, X.-L. Zhang, D.-W. Pang, J. Zhao, IAA stimulates pollen tube growth and mediates the modification of its wall composition and structure in Torenia fournieri. J. Exp. Bot. 59, 2529–2543 (2008).

26. F. Roda, G. M. Walter, R. Nipper, D. Ortiz-Barrientos, Genomic clustering of adaptive loci during parallel evolution of an Australian wildflower. Mol. Ecol. 26, 3687–3699 (2017).

27. M. E. James, H. Arenas-Castro, J. S. Groh, J. Engelstädter, D. Ortiz-Barrientos, Highly replicated evolution of parapatric ecotypes. bioRxiv 10.1101/2020.02.05.936401, 2020.2002.2005.936401 (2020).

28. J. Friml, Auxin transport—shaping the plant. Curr. Opin. Plant Biol. 6, 7–12 (2003).

29. D. Lopez, K. Tocquard, J.-S. Venisse, V. Legué, P. Roeckel-Drevet, Gravity sensing, a largely misunderstood trigger of plant orientated growth. Front. Plant Sci. 5 (2014).

30. D. Sang et al., Strigolactones regulate rice tiller angle by attenuating shoot gravitropism through inhibiting auxin biosynthesis. Proc. Natl. Acad. Sci. USA 111, 11199–11204 (2014).

31. M. C. Melo, A. Grealy, B. Brittain, G. M. Walter, D. Ortiz-Barrientos, Strong extrinsic reproductive isolation between parapatric populations of an Australian groundsel. New Phytol. 203, 323–334 (2014).

32. G. M. Walter et al., Diversification across a heterogeneous landscape. Evolution 70, 1979–1992 (2016).

33. G. M. Walter, M. J. Wilkinson, J. D. Aguirre, M. W. Blows, D. Ortiz-Barrientos, Environmentally induced development costs underlie fitness tradeoffs. Ecology 99, 1391–1401 (2018).

34. T. J. Richards, G. M. Walter, K. McGuigan, D. Ortiz-Barrientos, Divergent natural selection drives the evolution of reproductive isolation in an Australian wildflower. Evolution 70, 1993–2003 (2016).

35. F. Roda et al., Genomic evidence for the parallel evolution of coastal forms in the Senecio lautus complex. Mol. Ecol. 22, 2941–2952 (2013).

36. G. M. Walter et al., Loss of ecologically important genetic variation in late generation hybrids reveals links between adaptation and speciation. Evolution Letters 4, 302–316 (2020).

37. L. Fishman, J. H. Willis, Evidence for Dobzhansky-Muller incompatibilites contributing to the sterility of hybrids between Mimulus guttatus and M. nasutus. Evolution 55, 1932–1942 (2001).

38. R. B. Stelkens, C. Schmid, O. Seehausen, Hybrid breakdown in cichlid fish. PLOS One 10, e0127207 (2015).

39. M. Yamamoto, K. T. Yamamoto, Differential effects of 1-naphthaleneacetic acid, indole-3-acetic acid and 2,4-dichlorophenoxyacetic acid on the gravitropic response of roots in an auxin-resistant mutant of Arabidopsis, auxl. Plant Cell Physiol. 39, 660–664 (1998).

40. I. Ottenschläger et al., Gravity-regulated differential auxin transport from columella to lateral root cap cells. Proc. Natl. Acad. Sci. USA 100, 2987–2991 (2003).

41. E. B. Blancaflor, Regulation of plant gravity sensing and signaling by the actin cytoskeleton. Am. J. Bot. 100, 143–152 (2013).

42. D. B. Lowry, D. Popovic, D. J. Brennan, L. M. Holeski, Mechanisms of a locally adaptive shift in allocation among growth, reproduction, and herbivore resistance in Mimulus guttatus*. Evolution 73, 1168–1181 (2019).

43. L. Barboza et al., Arabidopsis semidwarfs evolved from independent mutations in GA20ox1, ortholog to green revolution dwarf alleles in rice and barley. Proc. Natl. Acad. Sci. USA 110, 15818–15823 (2013).

44. P. Ranocha et al., Walls are thin 1 (WAT1), an Arabidopsis homolog of Medicago truncatula NODULIN21, is a tonoplast-localized protein required for secondary wall formation in fibers. The Plant Journal 63, 469–483 (2010).

45. P. Ranocha et al., Arabidopsis WAT1 is a vacuolar auxin transport facilitator required for auxin homoeostasis. Nature Communications 4, 2625 (2013).

46. S. Promchuea, Y. Zhu, Z. Chen, J. Zhang, Z. Gong, ARF2 coordinates with PLETHORAs and PINs to orchestrate ABA-mediated root meristem activity in Arabidopsis. J. Integr. Plant Biol. 59, 30–43 (2017).

47. Dai, K.-i. Hayashi, H. Nozaki, Y. Cheng, Y. Zhao, Genetic and chemical analyses of the action mechanisms of sirtinol in Arabidopsis. Proceedings of the National Academy of Sciences of the United States of America 102, 3129 (2005).

48. M. Sagi, C. Scazzocchio, R. Fluhr, The absence of molybdenum cofactor sulfuration is the primary cause of the flacca phenotype in tomato plants. The Plant Journal 31, 305–317 (2002).

49. H. Wu, Y. Shen, Y. Hu, S. Tan, Z. Lin, A phytocyanin-related early nodulin-like gene, BcBCP1, cloned from Boea crassifolia enhances osmotic tolerance in transgenic tobacco. J. Plant Physiol. 168, 935–943 (2011).

50. H. Ma, H. Zhao, Z. Liu, J. Zhao, The phytocyanin gene family in rice (Oryza sativa L.): Genome-wide identification, classification and transcriptional analysis. PLOS One 6, e25184 (2011).

51. J. Garcia de la Garma et al., New insights into plant salt acclimation: the roles of vesicle trafficking and reactive oxygen species signalling in mitochondria and the endomembrane system. New Phytol. 205, 216–239 (2015).

52. Y. Hou et al., Maternal ENODLs are required for pollen tube reception in Arabidopsis. Curr. Biol. 26, 2343–2350 (2016).

53. H.-H. Wang et al., Close arrangement of CARK3 and PMEIL affects ABA-mediated pollen sterility in Arabidopsis thaliana. Plant, Cell Environ. 43, 2699–2711 (2020).

54. A. Sicard et al., Divergent sorting of a balanced ancestral polymorphism underlies the establishment of gene-flow barriers in Capsella. Nature Communications 6, 7960 (2015).

55. A. L. Sweigart, L. E. Flagel, Evidence of natural selection acting on a polymorphic hybrid incompatibility locus in Mimulus. Genetics 199, 543–554 (2015).

56. W. Beeftink, J. Rozema, A. Huiskes, Ecology of coastal vegetation (Springer, 1985), vol. 6.

57. T. Auld, D. Morrison, Genetic determination of erect and prostrate growth habit in five shrubs from windswept headlands in the Sydney region. Aust. J. Bot. 40, 1–11 (1992).

58. D. Morrison, A. Rupp, Patterns of morphological variation within Acacia suaveolens (Mimosaceae). Aust. Syst. Bot., 1013–1027 (1995).

59. G. M. Crutsinger, S. Y. Strauss, J. a. Rudgers, Genetic variation within a dominant shrub species determines plant species colonization in a coastal dune ecosystem. Ecology 91, 1237–1243 (2010).

60. T. Li et al., Calcium signals are necessary to establish auxin transporter polarity in a plant stem cell niche. Nature Communications 10, 726 (2019).

61. V. Naser, E. Shani, Auxin response under osmotic stress. Plant Mol. Biol. 91, 661–672 (2016).

62. T. van den Berg, R. A. Korver, C. Testerink, K. H. ten Tusscher, Modeling halotropism: a key role for root tip architecture and reflux loop remodeling in redistributing auxin. Development 143, 3350–3362 (2016).

63. C. S. Galvan-Ampudia et al., Halotropism is a response of plant roots to avoid a saline environment. Curr. Biol. 23, 2044–2050 (2013).

64. D. B. Lowry, J. H. Willis, A widespread chromosomal inversion polymorphism contributes to a major life-history transition, local adaptation, and reproductive isolation. PLoS Biol. 8, e1000500 (2010).

65. D. B. Lowry, M. C. Hall, D. E. Salt, J. H. Willis, Genetic and physiological basis of adaptive salt tolerance divergence between coastal and inland Mimulus guttatus. New Phytol. 183, 776–788 (2009).

66. Y. Yang, U. Z. Hammes, C. G. Taylor, D. P. Schachtman, E. Nielsen, High-affinity auxin transport by the AUX1 influx carrier protein. Curr. Biol. 16, 1123–1127 (2006).

67. C. A. Schneider, W. S. Rasband, K. W. Eliceiri, NIH Image to ImageJ: 25 years of image analysis. Nat. Methods 9, 671–675 (2012).

68. G. Rigo et al., Inactivation of plasma membrane-localized CDPK-RELATED KINASE5 decelerates PIN2 exocytosis and root gravitropic response in Arabidopsis. The Plant Cell 25, 1592–1608 (2013).

69. A. M. Rashotte, S. R. Brady, R. C. Reed, S. J. Ante, G. K. Muday, Basipetal auxin transport is required for gravitropism in roots of Arabidopsis. Plant Physiol. 122, 481–490 (2000).

70. R Core Team, R: A language and environment for statistical computing. R Foundation for Statistical Computing, Vienna, Austria (2013).

71. M. Lynch, B. Walsh, Genetics and analysis of quantitative traits (Sinaeur Associates Inc, Sunderland, U.S.A, 1998).

72. M. V. Rockman, L. Kruglyak, Breeding designs for recombinant inbred advanced intercross lines. Genetics 179, 1069–1078 (2008).

73. C. J. Geyer, S. Wagenius, R. G. Shaw, Aster models for life history analysis. Biometrika 94, 415–426 (2007).

74. R. G. Shaw, C. J. Geyer, S. Wagenius, H. H. Hangelbroek, J. R. Etterson, Unifying life-history analyses for inference of fitness and population growth. The American Naturalist 172, E35–E47 (2008).

75. J. D. Clarke, Cetyltrimethyl ammonium bromide (CTAB) DNA miniprep for plant DNA isolation. Cold Spring Harb. Protoc. 2009, ppdb.prot5177 (2009).

76. H. Liu (2015) Developing genomic resources for an emerging ecological model species Senecio lautus. in School of Biological Sciences (The University of Queensland), p 274.

77. N. A. Baird et al., Rapid SNP discovery and genetic mapping using sequenced RAD markers. PLOS One 3, 1–7 (2008).

78. B. Langmead, C. Trapnell, M. Pop, S. L. Salzberg, Ultrafast and memory-efficient alignment of short DNA sequences to the human genome. Genome Biology 10, R25 (2009).

79. H. Li et al., The sequence alignment/map format and SAMtools. Bioinformatics 25, 2078–2079 (2009).

80. S. F. Altschul, W. Gish, W. Miller, E. W. Myers, D. J. Lipman, Basic local alignment search tool. J. Mol. Biol. 215, 403–410 (1990).

